# Mapping phosphorylation post-translational modifications along single peptides with nanopores

**DOI:** 10.1101/2022.11.11.516163

**Authors:** Ian C. Nova, Justas Ritmejeris, Henry Brinkerhoff, Theo J. R. Koenig, Jens H. Gundlach, Cees Dekker

**Affiliations:** Department of Bionanoscience, Kavli Institute of Nanoscience Delft, Delft University of Technology, Delft, Netherlands; Department of Physics, University of Washington, Seattle, WA, USA

## Abstract

Current methods to detect post-translational modifications (PTMs) of proteins, such as phosphate groups, cannot measure single molecules and often cannot differentiate between closely spaced phosphorylation sites. Using a nanopore sequencing approach, we here report detection of PTMs at the single-molecule level on immunopeptide sequences with cancer-associated phosphate variants. We reliably discriminate peptide sequences with one or two closely spaced phosphates with 95% accuracy for individual reads of single molecules.

---

Post-translational modifications (PTMs) play crucial roles in protein function and cell fate. Most PTMs involve attachment of a small chemical group (phosphoryl, acetyl, glycosyl, etc.) to amino acids, which greatly expands the proteome. Mass spectrometry is the principal technique to detect PTMs, but this method requires significant sample input (typically >10^9^ copies) and often struggles to identify the correct position of a PTM between multiple candidate sites^1^. Improved detection of protein phosphorylation, the most frequent PTM,^2^ is of particular interest, as dysregulation of phosphorylation pathways is linked to many diseases including cancers, Parkinson’s, Alzheimer’s, and heart disease.^3^ Specifically, certain phosphorylation patterns on immunopeptides, which are naturally digested protein products on the cell surface for immune cell recognition, have been directly linked to cancer cells, making these immunopeptide variants promising neoantigens (cancer-specific antigens) for targeted immunotherapy or cancer screening.^4^ Nanopore techniques, where the change in ion current is measured as a single molecule passes through a nanopore in a membrane, have shown promise for PTM detection.^5–15^ However, these approaches, which measure brief transient blockades, have so far lacked high accuracy in variant identification for single molecules.

Here, we apply a recently introduced nanopore single-protein scanning method^16–18^ to PTM detection and demonstrate its capabilities to detect and discriminate single phosphate groups within individual peptides. In this approach,^16^ a peptide of interest (up to ~25 amino acids) is chemically linked to a DNA oligonucleotide, creating a peptide-oligonucleotide conjugate (POC) which is slowly translocated in a stepwise manner through a nanopore (MspA^19^) using a DNA motor enzyme (He1308 helicase^20^), as in nanopore DNA sequencing.^21–24^ Previously,^16^ individual amino acid substitutions on single peptides were discriminated with high accuracy, but the peptide sequence tested was atypical, with a near-uniform negatively charged chain of aspartate and glutamate residues to induce electrophoretic insertion of the POC into the nanopore. To test biologically relevant peptides with various charges, we chemically linked a second DNA oligo (the ‘threading DNA’) to the other end of the peptide^18^ (Fig. 1a). This DNA electrophoretically threads the POC into and through the nanopore where it is subsequently pulled back out of the pore in ~0.3 nm steps by the helicase, slowly scanning the peptide across the sensing region of the pore, i.e., the constriction point of MspA which controls most of the ion current (Fig. 1b). Fig. 1c depicts a typical ion current trace from a single translocation event of a POC containing an 10 amino acid peptide. The first part of the trace reads the template DNA section which corresponds well with the predicted pattern from nanopore DNA sequencing^24^ (Fig. 1d), whereas the second part contains the linker and peptide signal of interest.

**Fig. 1:**
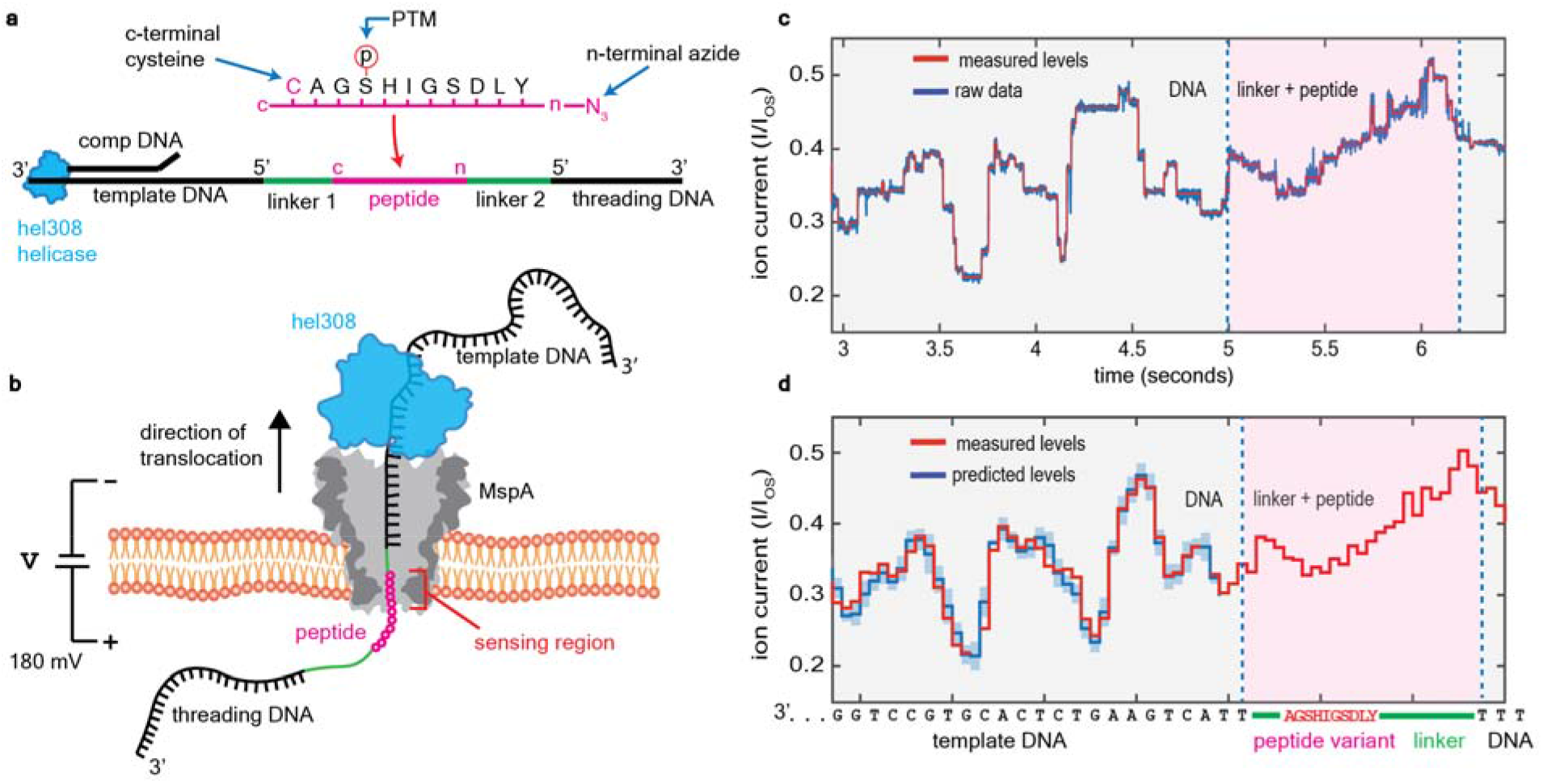
Nanopore PTM detection experimental schematic and data workflow. **a,** Schematic of the peptide-oligo construct (POC). A (phosphorylated) immunopeptide (pink) is linked by its c-terminus to the 5’ end of a DNA template (linker 1, cysteine-maleimide bond), while its n-terminus is linked to the 5’ end of the threading DNA (linker 2, azide-DBCO bond). Hel308 helicase loads onto the ssDNA/dsDNA junction made by a complementary oligo (comp DNA) that is annealed to the template DNA. **b,** Schematic of POC reading. A MspA nanopore (grey) is embedded in a lipid bilayer. Applied voltage (180 mV) drives a current of K^+^ and Cl^-^ ions through the nanopore. The threading DNA is electrophoretically driven into and through the nanopore, translocating the POC, stripping off the comp DNA, and docking the Hel308 onto the rim of MspA. As Hel308 steps along the template DNA, the POC is pulled up through the pore in ~0.33 nm increments, thereby pulling residues through the narrowest portion of MspA (sensing region) where they modulate the ion current. **c,** Ion current trace of a typical POC reading event for βCAT. Ion currents (I) are normalized to the unblocked open-state pore current (I_OS_). Measured levels (red) are determined using a level-finding algorithm. After reading the template DNA, linker 1 enters the sensing region (at 5 s), followed by peptide, linker 2, and the start of the threading DNA. **d,** Consensus sequence of ion current steps (red), which for the DNA section is closely matched by the predicted DNA sequence (blue). Error bars in the measured ion current levels are errors in the mean value, often too small to see. Error bars in the prediction are standard deviations of the ion current levels that were used to build the predictive map.^27^

We found that this approach allows extremely sensitive measurements that can clearly distinguish peptides with or without a single PTM. We measured POCs containing the immunopeptide BCAR3 (Fig. 2a), a promising neoantigen for immunotherapy. We compared BCAR3 (with sequence LKEPTRDMI, written c to n terminus) and its phosphate-PTM-containing variant pBCAR3 where a single threonine residue was phosphorylated (LKEP[pT]RDMI). Consensus ion current patterns were determined by aligning and averaging n=40 reads of each variant, see Fig. 2b. The addition of phosphothreonine (pT), a single small PTM of only 5 atoms, produced a dramatic change to the current pattern. Specifically, the pattern for the phosphate-containing variant was consistent with unphosphorylated BCAR3 until pT entered the sensing region, whereupon the current increased significantly by up to 25% for 13 steps, until the current returned to match for the rest of the remaining steps. These data clearly show that a single PTM can be well distinguished in nanopore reading.

**Fig. 2:**
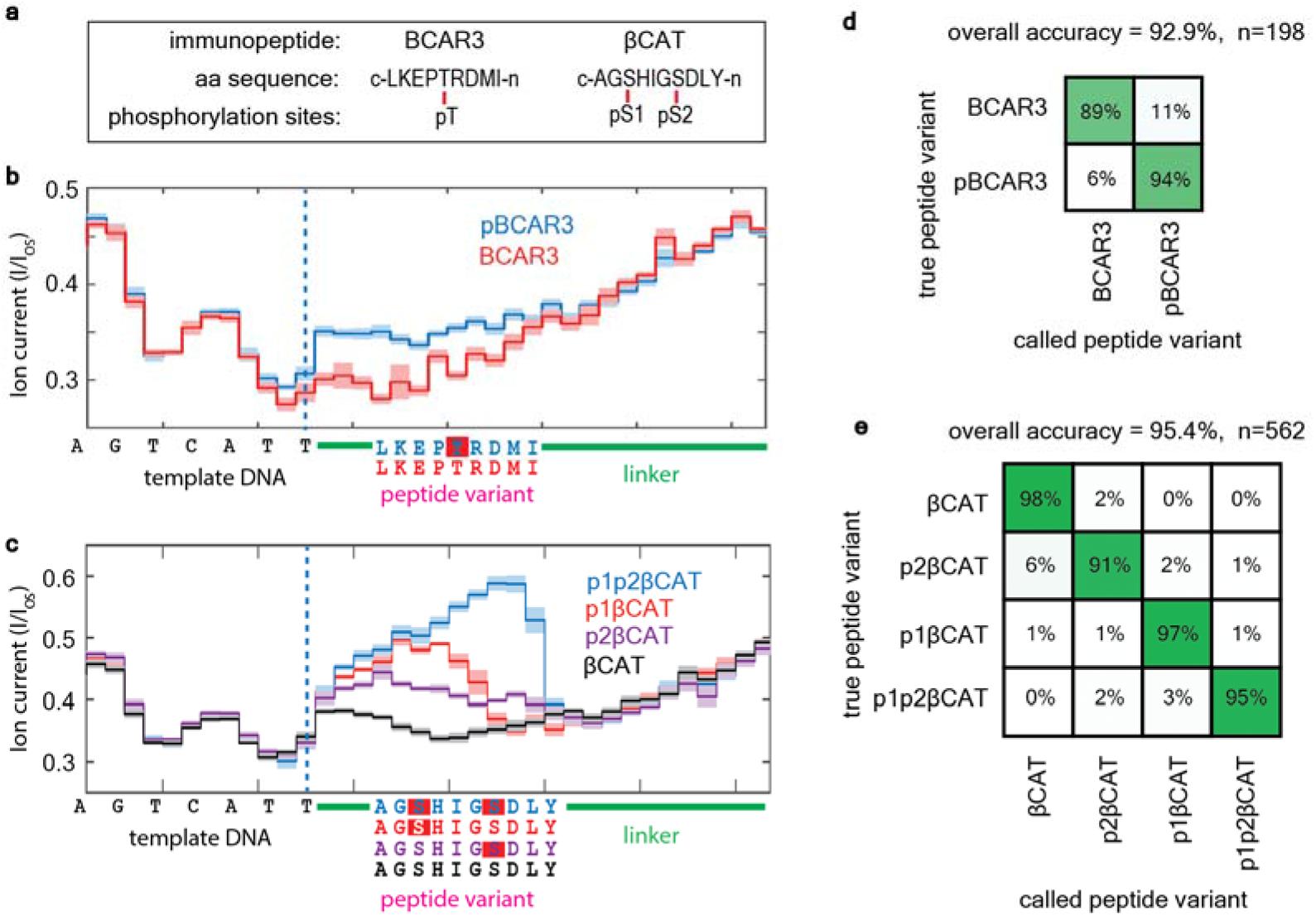
Discrimination of immunopeptide PTM phosphate variants. **a,** Immunopeptides with amino acid sequences and phosphorylation sites. BCAR3 contains a single phosphorylation site at a threonine residue (pT). βCAT contains two serine phosphorylation sites (termed pS1 and pS2) separated by 3 amino acids. Phosphopeptide variants studied were BCAR3, pBCAR3 (with pT), βCAT, p1βCAT (with pS1), p2βCAT (with pS2) and p1p2βCAT (with both pS1 and pS2). **b,** Consensus ion current patterns for BCAR3 (and for the PTM variant pBCAR3 (N=40 reads for each trace). Dashed line marks the end of the template DNA in the sensing region. **c,** Consensus ion current patterns for βCAT and its phosphopeptide variants (N=40 reads for each trace). **d,** Single-read blinded-variant-calling matrix for BCAR3 variants yielding an overall variant-calling accuracy of 93%. **e,** Same for βCAT variants, yielding an overall variant-calling accuracy of 95%.

We next demonstrated the sensitivity of this method to discriminate between closely spaced PTMs along a peptide. We repeated the procedure to analyze another clinically relevant immunopeptide, βCAT (AGSHIGSDLY), that contains *two* phosphorylation sites, one at each serine (termed pS1 and pS2), at positions separated by three amino acids (Fig. 2a). We determined the current patterns for the unphosphorylated variant, both single-phosphoserine (pS) variants (p1βCAT containing pS1, AG[pS]HIGSDLY; and p2βcat containing pS2, AGSHIG[pS]DLY), and the double pS variant (p1p2βCAT containing both pS1 and pS2, AG[pS]HIG[pS]DLY) (Fig. 2c). All four βCAT variants produced a distinct ion current pattern that could clearly be discriminated from that of the other variants. Just like for pT (Fig. 2b), the addition of pS had the consistent effect of increasing the current. Notably, the magnitude of the increase and the number of steps that were affected varied between the two single phosphorylation sites (9 steps for p1βCAT and 12 steps for p2βCAT). For the double phosphopeptide (p1p2βCAT), the two phosphoserines combined to increase the current even more than with the two single variants, reaching large current values that exceeded the nonphosphorylated variant by up to 64% for 12 steps.

These differences in ion current patterns can be used to accurately identify the correct variant for individual reads of these immunopeptides – as can be quantified in a so-called confusion matrix. For 198 single reads of BCAR3 and its variant, we blindly determined the correct variant using a hidden Markov model with an accuracy of 93%, see Fig. 2d. For 562 reads of βCAT and its variants, we determined the correct variant with 95% accuracy, while individual variant-calling accuracies ranged between 91% (βCAT) and 98% (p2βCAT) (Fig. 2e). Overall, the single-read variant-calling accuracy was 95% for all of the measured phosphopeptides, highlighting the capabilities of this technique to reliably determine the correct PTM location on single molecules.

The heterogeneous charge profile of these peptides leads to variations in the POC polymer’s stretching as it is stepped through the pore. The constant *k*-mer reading frame^21^ that is commonly used in models of nanopore DNA sequencing is therefore inadequate to describe the influence of amino acid sequence on ion current patterns. We developed a physical model to better understand this behavior. For each of the four βCAT variants, we performed a Markov-chain Monte Carlo (MCMC) calculation (see Methods), where the POC was modeled as a freely jointed chain with units of varying charge (Fig. 3a), anchored at the top of the MspA pore by Hel308, and subject to ion-screened Coulomb forces between charges as well as to the applied electrostatic potential (Fig. 3b). By performing the MCMC calculation with each βCAT variant at 30 consecutive Hel308 steps, we tracked the movement of the POCs through the nanopore. Figure 3c depicts typical configurations found for p2βCAT at a selection of Hel308 steps, while Fig. 3d plots the corresponding mean *z*-location (vertical axis along the pore) of the pS2 PTM, calculated for every step. We find that after the template DNA is stepped through the sensing region, the linker/peptide polymer bunches within the pore, until the large negative charge (pS2) is held just below the nanopore constriction by the voltage drop. As stepping continues, the slack is gradually pulled out of the polymer and the phosphate is slowly pulled up into the pore constriction, reaching a critical point at which the charged phosphate quickly pops up into the pore vestibule and the polymer returns to a bunched slack configuration. This is illustrated in the trace of panel 3d where the pS2 PTM stalls at *z* ~ 8 nm, until it suddenly jumps up at step 19. While residing in the stalling position just below the pore, the negative phosphate group likely promotes the transit of K^+^ ions, thus increasing the nanopore current, as seen in the experimental data (Fig. 2c). The stalling and jumping behavior was consistently observed for all PTMs in all βCAT peptides, see Fig. 3e.

**Fig. 3.**
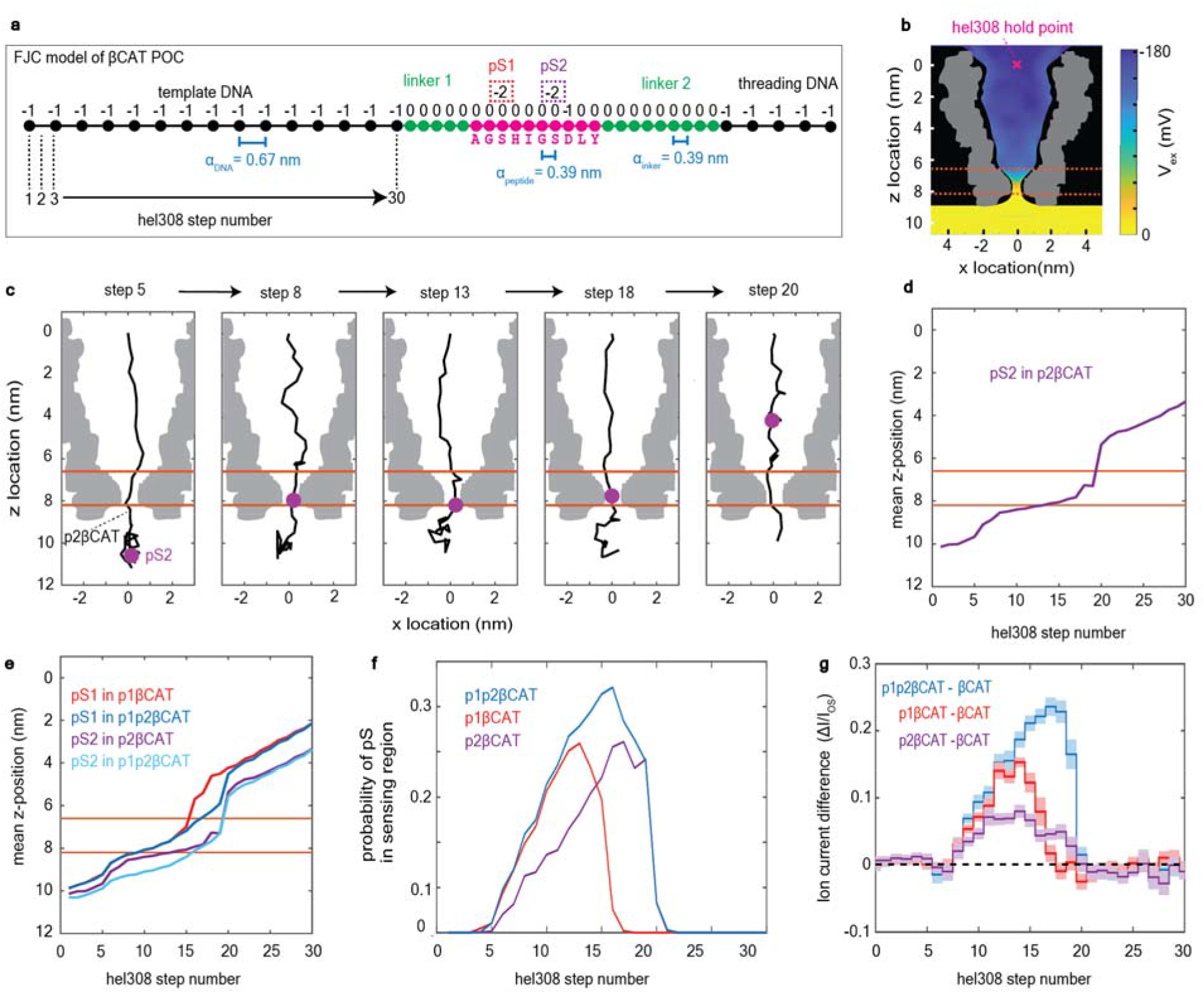
Markov-chain Monte Carlo (MCMC) calculations of phosphate-containing peptides. **a,** Freely-jointed chain (FJC) model of a POC containing peptide βCAT. Each unit in the polymer has an electrical charge and a typical distance α between residues. Phosphoserine PTMs pS1 and pS2 add a charge of −2 to that unit. **b,** Electric potential profile (color gradient) in the MspA pore.^30^ The POC was confined within the physical boundaries of MspA (black) and anchored at the Hel308 hold point (pink x). The volumetric map of the MspA cross-section is shown in grey. **c,** Snapshots of the polymer configuration within MspA from MCMC calculations for the p2βCAT POC at five Hel308 steps. The phosphoserine residue (pS2) (purple) is observed to move through the pore in a nonlinear fashion. Note that the POC polymer gets stretched towards Hel308 step 19, after which the PTM moves into the pore lumen and the polymer relaxes. Orange lines indicate the sensing region of MspA. **d,** Mean z-location of the pS2 PTM versus Hel308 step number. **e,** Same for all PTMs in the peptide variants. **f,** Probability that a pS occupied the sensing region for various βCAT PTM variants versus Hel308 step. **g,** Experimentally measured ion currents for βCAT phosphopeptide variants where the ion current measured for the non-PTM βCAT was subtracted (from data in Fig. 2c.).

As a proxy for the ion current patterns, we extracted the percentage of time that a phosphate PTM was present in the sensing region (defined as 6.6nm < z < 8.2nm, see Methods) at each step in the simulations (Fig. 3f). The results display an excellent correspondence with the experimental ion current differences (Fig. 3g), capturing the wider region of influence for pS2 in p2βCAT compared to pS1 in p1βCAT (12 steps vs. 9 steps). In addition, the combined effect of pS1 and pS2 in p1p2βCAT influencing the same region as pS2 on its own in p2βCAT (12 total steps for both, Fig. 3g) is well represented by the model (Fig. 3f). Overall, this model provides a starting point for understanding how the charge distribution along peptides relates to ion current traces.

Our data provide the first demonstration of a technique that can accurately differentiate between single-molecule phosphopeptide variants upon controllably drawing the peptide along the sensing region of a nanopore. Importantly, the technique can clearly distinguish phosphopeptides with phosphates that are separated by only a few amino acids (3 in our example), where mass spectrometry faces particular difficulties, and it does not require chemical labelling of the PTM as in other single-molecule proteomics methods.^25^ The variant-calling accuracy that we realized was already very high (95%) in single reads, but can, if desired, be further improved upon using the rereading capability of our nanopore scanning approach.^16^ The results presented here can logically be extended to differentiate between phosphopeptide variants of any peptide sequence. In addition, detection of other PTM types (acetylation, hydroxylation, methylation, etc.) can likely be achieved with an identical approach, as long as the PTM is not too bulky to translocate through MspA. Further developments may move from using synthetic peptides to natural peptides harvested from a biological sample, increasing throughput using arrays of nanopores in parallel, and developing robust methods to attach DNA to the n and c terminus of peptides without a priori modifications. This demonstration of single PTM detection within individual peptides presents a new tool for phosphorylation investigation, enabling measurements currently unachievable with other proteomics tools.

## Methods

### Construction of peptide-oligo conjugates

Peptides (sequences in Supplementary Table 1) were purchased from Life Technologies and diluted to 10 μM in degassed PBS buffer. DNA oligos (sequences in Supplementary Table 2) were purchased from Biomers and diluted to 100 μM in degassed PBS buffer. Orthogonal click-chemistry reactions were used to attach a C terminal cysteine on the peptide to a 5’ maleimide on the template DNA, and to attach a N terminal azide on the peptide to a 5’ DBCO on the threading DNA.

The cysteine-maleimide reaction and the DBCO-azide copper-free click reaction were performed in one pot. Peptides and DNAs were mixed at a ratio of 1:2:6 (peptide: threading DNA: template DNA) at a concentration of 7 μM peptide in PBS and were incubated overnight (20h) at +4C under argon gas. Supplementary Fig. 1 depicts the chemical structure of the full POC. The mixture was purified using DynaBeads strep-biotin polyA cleanup, ensuring that only constructs containing threading DNA (poly T) remained. The resulting product was estimated to be at a final concentration of 12.5 μM based on the maximum binding capacity of the beads. Comp DNA was added at a concentration of 15 μM and annealed to template DNA.

### Nanopore experimental methods

Nanopore measurements were conducted as in Brinkerhoff et al.^16^ and previous studies^19–24^ with a few notable differences. DPhPC lipids purchased from Avanti were used to paint bilayers on ~10 μM Teflon apertures in custom U-tube experimental devices. MspA mutant M2-NNN^19^ was purified by Genscript. All experiments were conducted at 37±1 °C with 1 mM ATP, 400 mM KCl, 10 mM MgCl2, 10 mM HEPES pH 8.00 ± 0.05 in the *cis* well and 400 mM KCl, 10 mM HEPES pH 8.00 ± 0.05 in the *trans* well. Hel308 was added to *cis* well to a final concentration of 50 nM. POCs were added to a final concentration of 5 nM. Hel308 used in this study is from Thermococcus gammatolerans (accession number WP_015858487.1) and was cloned into the pET-28b(+) vector plasmid at Ndel/NotI sites by Genscript. Ion current data was acquired at 50 kHz sampling frequency using an Axopatch 200B patch clamp amplifier and filtered with 10 kHz 4-pole Bessel filter. Applied voltage was set to 180 mV for all experiments and controlled by a National Instruments X series DAQ and operated with custom LabVIEW software. Using these methods, many ion current reads (termed ‘events’) were gathered for each of the 6 POCs used in this study.

### Data analysis

All data analysis was performed in Matlab. Custom Matlab software as described in detail in Brinkerhoff et al.^16^ and briefly below, was used for data preprocessing, reduction, filtering, alignment, and variant identification:

#### Event selection and filtering

Data was downsampled to 5 kHz and potential events were identified using a thresholding algorithm based on the unblocked pore current as in previous work.^16,19–24^ Events were then selected by eye for further analysis. Occasionally, Hel308 fell off of the DNA before the end of the template strand. Therefore, we selected events that included steps for both template DNA and the entire peptide region into the second linker (see Supplementary Fig. 2 for example events that fit selection criteria).

#### Level finding and filtering

In order to determine the transition points between Hel308 steps in the data, we used a change point algorithm as described in previous work^19^ and originally developed in Wiggins et al.^28^ A sample of the typical behavior of this change point ‘level finder’ across an entire event is shown in Supplementary Fig. 3. These measured levels were further filtered, first by excluding any levels outside the bounds of expected current values (I/I_OS_ < 0.15 or I/I_OS_ > 0.7). Levels outside of these bounds correspond to noise spikes or mid-event gating of MspA pore. We next applied a recombination filter, as described in Noakes et al.,^27^ which identifies helicase backsteps and eliminates repeated levels from the trace. We delineated each event by eye by noting the position of the end of the DNA template in the measured levels, creating a DNA section (before this position) and peptide section (after this position, including both linker 1 and peptide and linker 2) for each event.

#### Reread removal

In our previous study,^16^ it was determined that at high Hel308 concentration, a string of multiple Hel308 enzymes can be loaded onto the DNA template strand during translocation. After the first Hel308 reaches the end of the DNA template and dissociates, the POC falls back through the pore until the next Hel308 sits on the rim of the pore and continues stepping. This produces a “reread” of the polymer, where the reread usually includes the final ~16 steps (equal to the footprint of one Hel 308 enzyme, 8 DNA bases). In the experiments presented here, the rereads did not include the variable region within the peptide but merely included levels corresponding to linker 2 within the pore (see Supplementary Fig. 4), while rereading of the relevant region can be engineered by a different linker design. In the current study, rereads were removed for subsequent analysis. Supplementary Fig. 4 depicts a typical rereading pattern and sections of removal.

#### DNA level prediction

The current pattern for the template DNA sequence was predicted using an empirically derived 6-mer map,^28^ where each 6 base sequence was given two ion current states, corresponding to the two substeps of Hel308 helicase (“pre-” and “post” steps) per DNA base. The ion currents in the map are the mean of the set of ion currents assigned to each state, and the uncertainty in the ion current is the standard deviation in that set of ion currents. The prediction matched well with the experimentally measured DNA levels in this study (Fig. 1d).

#### Initial consensus generation

For the peptide section of the measured ion current, the ion current patterns had to be experimentally determined for each POC variant, as no predictive map exists for peptides or other polymers that are not DNA or RNA. In order to determine the ion current patterns for the linker and peptide regions of each POC (Fig. 2), we determined an initial “best guess” of the ion current pattern. A selection of typical reads (n= 6-10) of each construct were compared by eye in order to determine the unique ion current states and place them in the appropriate order. This process eliminated Hel308 backsteps, repeated levels, and spurious states that are not representative of typical reads. These initial consensuses included the last 15 steps of the DNA template section and 28 steps after the DNA template (where the helicase typically fell off of the DNA).

Reads were then calibrated, applying a scaling factor *m* to the measured ion current to account for slight variations in buffer salt concentration due to evaporation. Determination of the scaling factor was done as in Brinkerhoff et al.,^16^ where the maximum likelihood estimator for *m* that limited the error between reads was calculated for each read. After applying the scales, a mean and standard deviation value was calculated for each position in the consensus. Next, a second round of calibration was applied to the mean consensus values in order to ensure cross-calibration consistency between consensus of the different POC variants. We calculated a scale and offset factor by performing a single-polynomial fit of the first 15 steps of each initial consensus (corresponding to the DNA section) to the last 15 levels of the predicted ion current pattern for the DNA template sequence. This ensured that all of the consensus patterns were calibrated to the same reference.

#### Final consensus generation

These calibrated initial consensuses were then used as initial guesses for a customized Baum-Welch algorithm, a type of expectation maximization for the hidden Markov model. This algorithm was performed identically as in Brinkerhoff et al.^16^ and described fully in Noakes et al.^27^ We randomly chose 40 events of each POC variant for the EM algorithm. In order to calibrate these events, we performed a hidden-Markov-model alignment of the levels in the DNA section of each event to the template DNA prediction over a range of scale factors (*m*= 0.8 to 1.2 with increments of 0.01) and calculated a likelihood score for each *m* value. We chose the *m* value that produced the highest alignment score. We then applied this event specific scale factor to the associated measured levels from the peptide section of the same event. The expectation maximization algorithm was then used to generate a final consensus for each POC variant, using this set of calibrated events of each variant and the initial consensus as a seed for the algorithm. Using this procedure, we obtained six final consensus ion current patterns (one for each of the immunopeptide variants used in this study). Fig. 2b and 2c depict the final consensus ion current patterns for each immunopeptide variant.

#### Variant identification

All filtered events that were not included in the initial or final consensus were used for variant identification. We calibrated each set of peptide levels using the DNA section alignment as previously. For each set of now calibrated peptide section levels, we performed a hidden Markov model alignment to the final consensus for each variant. Events containing βCAT and its associated variants were separated from BCAR3 and its associated variant and only aligned to the set of variants of the appropriate immunopeptide. The alignment producing the maximum alignment score was chosen as “called variant” (Fig. 2d and 2e). Alignment accuracy for each variant was calculated as the percentage of correct calls compared to the total number of calls. The overall accuracy was calculated by calculating the percentage of correct calls for all variants divided by total calls.

### Simulation methods

Markov-chain Monte Carlo calculations^29^ were implemented in MATLAB. Degrees of freedom were encoded as polar and azimuthal angles between each polymer joint, with the first joint being fixed at the origin located at the top of MspA’s vestibule. Spacings between joints were chosen to be 0.67 nm for DNA and 0.39 nm for linker and peptide regions. Charges were assigned to each joint: +1e^-^ for each K or R residue, −1e^-^ for each D or E residue and for each DNA monomer, −2e^-^ for each phosphorylated residue, and 0 for all other joints (see Supplementary Fig. 5). The update distribution for both the polar and azimuthal angles was a normal distribution with mean 0 and standard deviation 0.05 radians.

The potential energy was calculated as the sum of (i) the interaction between the joint charges and a previously published electric potential map of MspA (Supplementary Fig. 6);^30^ (ii) the coulomb interaction between pairs of joint charges, screened with a Debye radius of 0.40 nm; and (iii) a hard wall excluding any polymer joints from the wall of MspA, defined using a cylindrically symmetric spline derived from the electric potential map.

The MCMC calculation was performed at different “enzyme steps” by removing monomers from the top part of the chain one by one, thereby shifting the entire sequence up through the constriction. At each step, the calculation started with a completely extended chain, with all polar and azimuthal angles set to 0, and the calculation was iterated 10^6^ times. We discarded the first 10^4^ samples at each step in order to allow for thermalization of the samples before inclusion in the calculated distributions (For a detailed description of the simulations see Supplementary text 1).

Fig. 3f was produced by computing the fraction of samples in which a phosphorylation lay in the region where the z-component of the electric field along the z-axis was greater than 1 k_B_T / nm • e^-^ (6.6nm < z < 8.2nm, Supplementary Fig. 7) Fig. 3d and 3e were produced by computing the mean position of the phosphorylated residue in the samples for each construct during each step.

## Acknowledgements

We thank A. Laszlo for discussions on the Markov-chain Monte Carlo calculations, J. van der Torre for help in troubleshooting POC construction, E. van der Sluis and A. Goutou for Hel308 purification, and A. Aksimentiev for discussions. The work was supported by funding from the Dutch Research Council (NWO) project NWO-I680 (SMPS) (CD); European Research Council Advanced Grant 883684 (CD); European Commission Marie Skłodowska-Curie Fellowship 897672 (HB); and NIH NHGRI project HG012544-01 (JG and CD).

## Contributions

H.B. and C.D. conceived of and designed the study. I.C.N, J.R., and T.J.R.K. performed nanopore experiments. J.R. established and troubleshooted the method for POC construction. I.C.N. performed computational analyses of the experimental data. H.B. performed the simulations. C.D. and J.H.G. supervised the work. I.C.N. wrote the initial manuscript draft and all authors contributed to the writing of the final manuscript.

