## Supplemental Materials for "Mapping phosphorylation post-translational modifications along single peptides with nanopores"

Nova et. al

*Delft University of Technology*

| Peptide | Sequence ( C -> N) |
|---|---|
| BCAR3 | CLKEPTRDMI - azidePEG4 |
| pBCAR3 | CLKEP[pT]RDMI - azidePEG4 |
| βCAT | CAGSHIGSDLY - azidePEG4 |
| p1βCAT | CAG[pS]HIGSDLY - azidePEG4 |
| p2βCAT | CAGSHIG[pS]DLY - azidePEG4 |
| p1p2βCAT | CAG[pS]HIG[pS]DLY - azidePEG4 |

**Table S1** Peptides used in this study. Each peptide was synthesized with an N-terminal azide group and C-terminal cysteine for click chemistry reactions.

| **Oligonucleotide** | **Sequence ( 5’ -> 3’)** |
|---|---|
| Template DNA | Maleimide-TTACTGAAGTCTCACGTGCCTGGTATATTAGCGTCCACTC TCACTATCGGATTCTACATCGGTCGTAGCC |
| Complement DNA | CCGATGTAGAATCCGATAGTGAGAGTTTTTTTTTTTTTTTTTTTT-Cholesterol |
| Threading DNA (T50) | DBCO-PEG4-TTTTTTTTTTTTTTTTTTTTTTTTTTTTTTTTTTTTTTTTTT TTTTTTTT |

**Table S2** Oligonucleotides used in this study. The complement DNA contains a cholesterol on the 3’ end which associates with the bilayer and increases the interaction rate of the final POC with the pore.

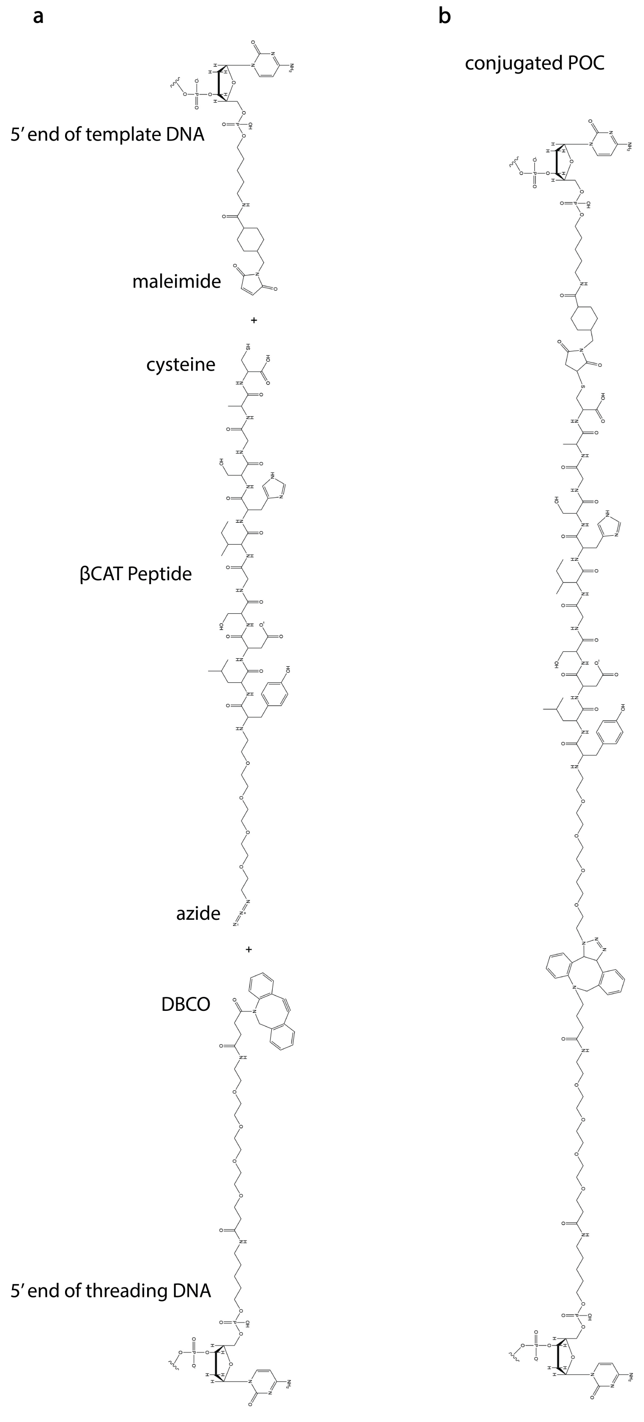


**Fig. S1** Chemical structure of the POC. A) Individual polymers being added for click chemistry reactions. The peptide $\beta$CAT is depicted. DNA sequences only include 1 base here in the structure but extend in the 3' direction. B) Full structure of conjugated product.

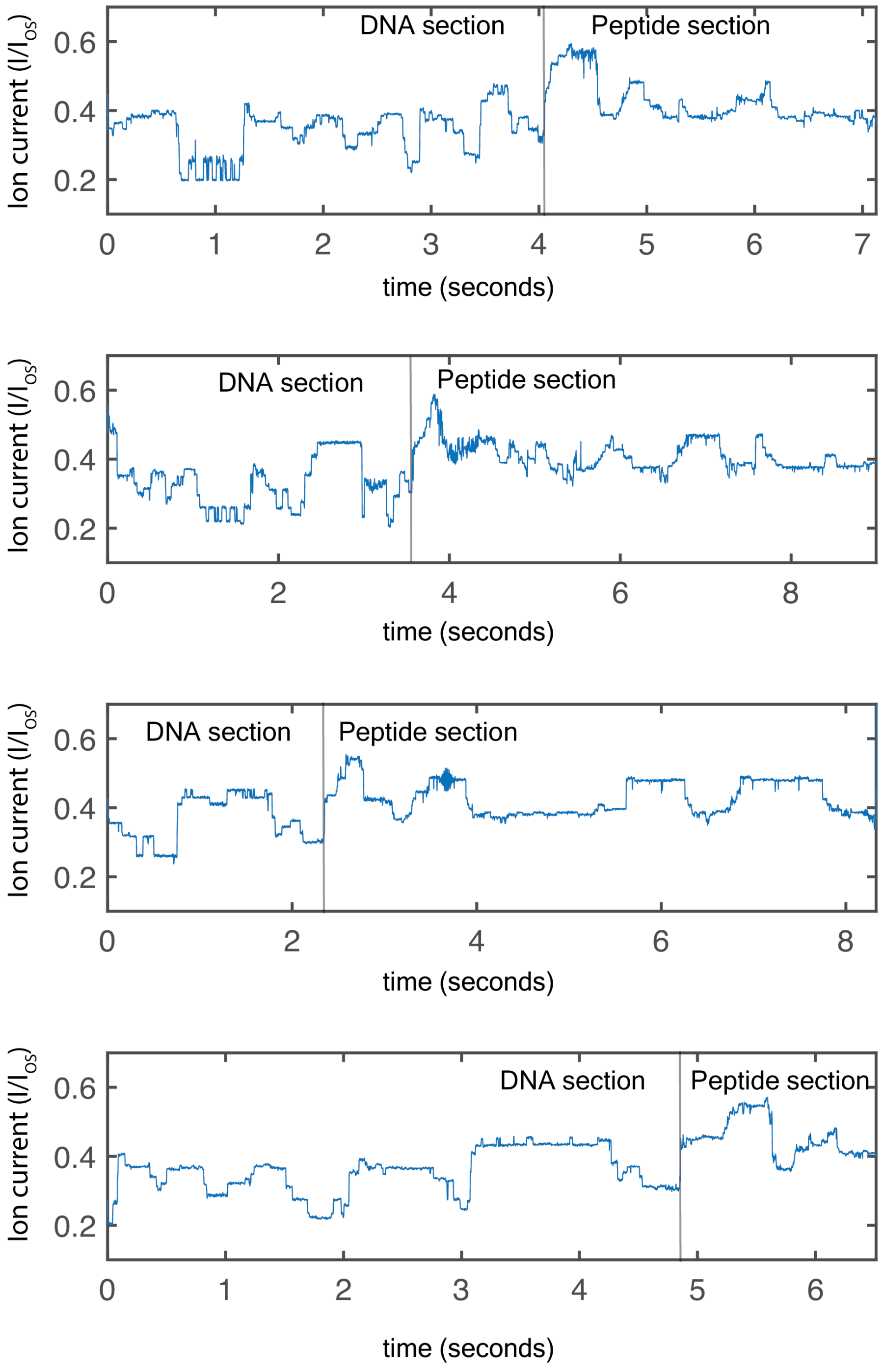


**Fig. S2** Example ion-current events containing p1p2$\beta$CAT POCs, downsampled to 500 Hz. The transition between the DNA section and peptide section is delineated by a vertical line. The peptide section includes linker1, peptide, linker 2 and the first base of the threading DNA. Events were selected that at least reached linker 2. The 4 events displayed here fit the criteria for selection for analysis.

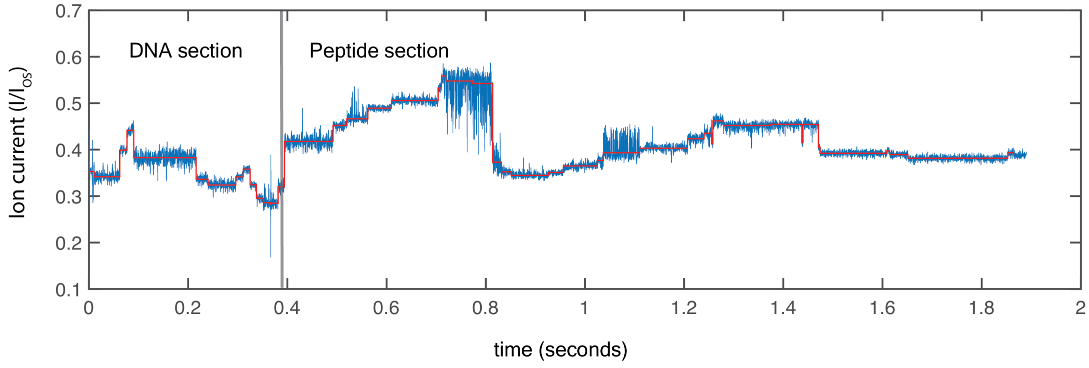


**Fig. S3** Typical level finding results. Typical p1p2$\beta$CAT read with measured levels. Raw data are in blue; fitted levels are in red. Data plotted at 5 kHz.

a

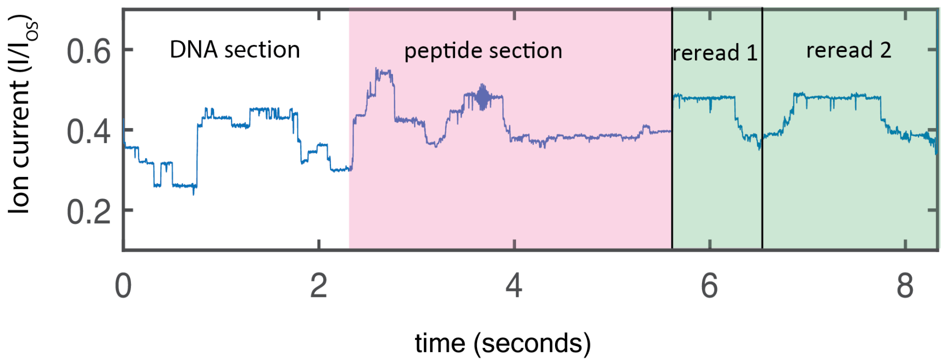


b

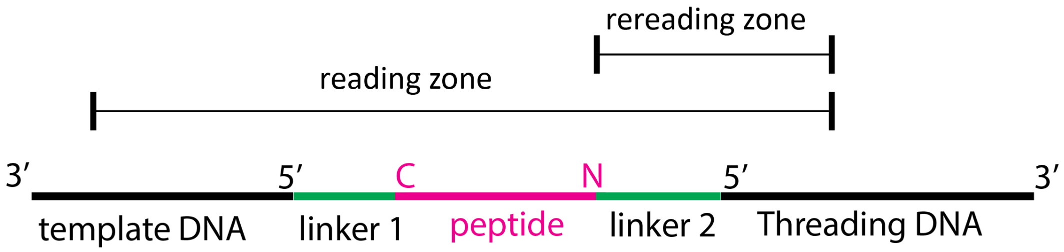


**Fig. S4** Rereading of POCs a. Example p1p2$\beta$CAT event (plotted at 500 Hz) showing the delineation of rereading. b. Schematic of POC with reading and rereading regions. The reading region corresponds to the section of polymer after the 5' end of the template strand that is sensed in the pore during the first read. The rereading zone is the area that is typically "reread" by the subsequent helicases bound to the POC.

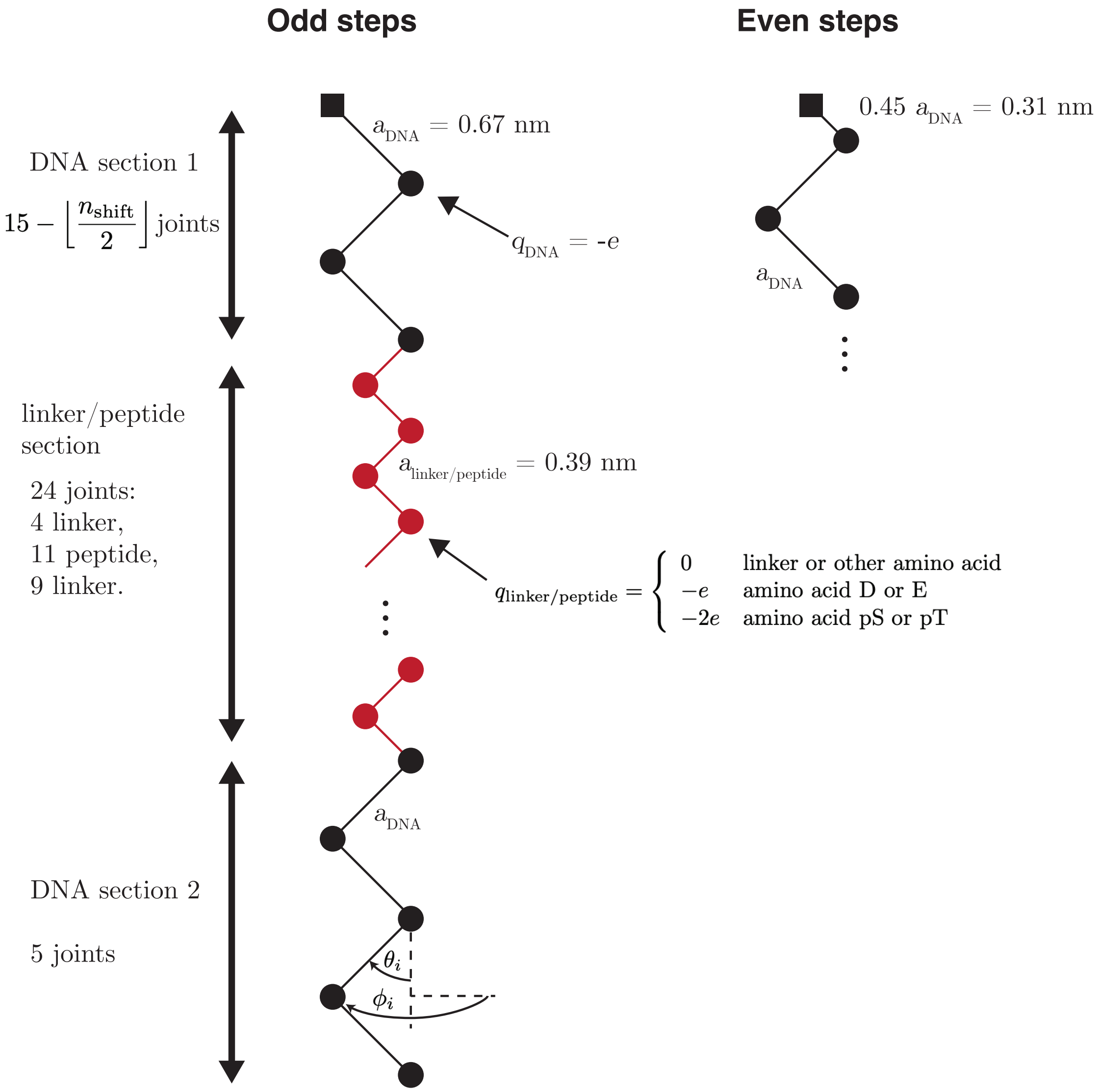


**Fig. S5** The freely-jointed chain model of the DNA-peptide-DNA construct. Hel308 takes two steps per DNA base (termed 'odd steps' and 'even steps'), which are modelled by shortening DNA section 1 by 0.36 nm for even steps and 0.31 nm for odd steps (based on the calculated step size of Hel308, (1)).

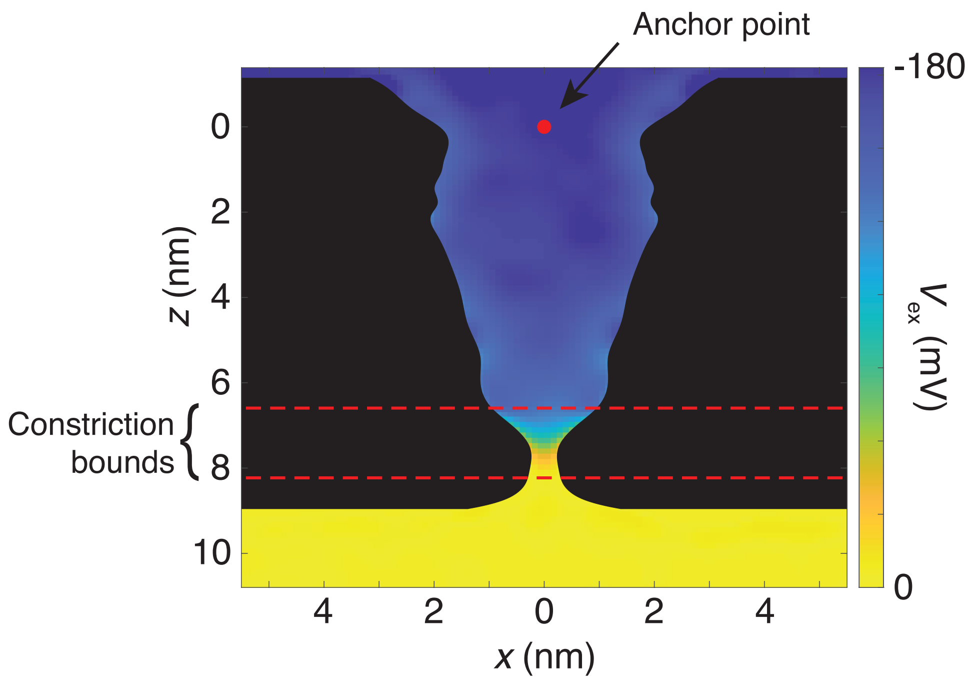


**Fig. S6** Diagram showing the electric potential $V_{\text{ex}}$ (colorbar), the pore radius $r_p(z)$ (black boundary), and the boundary of the high-field constriction as determined by threshholding the $z$-component of the electric field along the $z$-axis (red dashed lines; see Fig. S7). The endpoint of the freely-jointed chain is constrained to the origin (red dot).

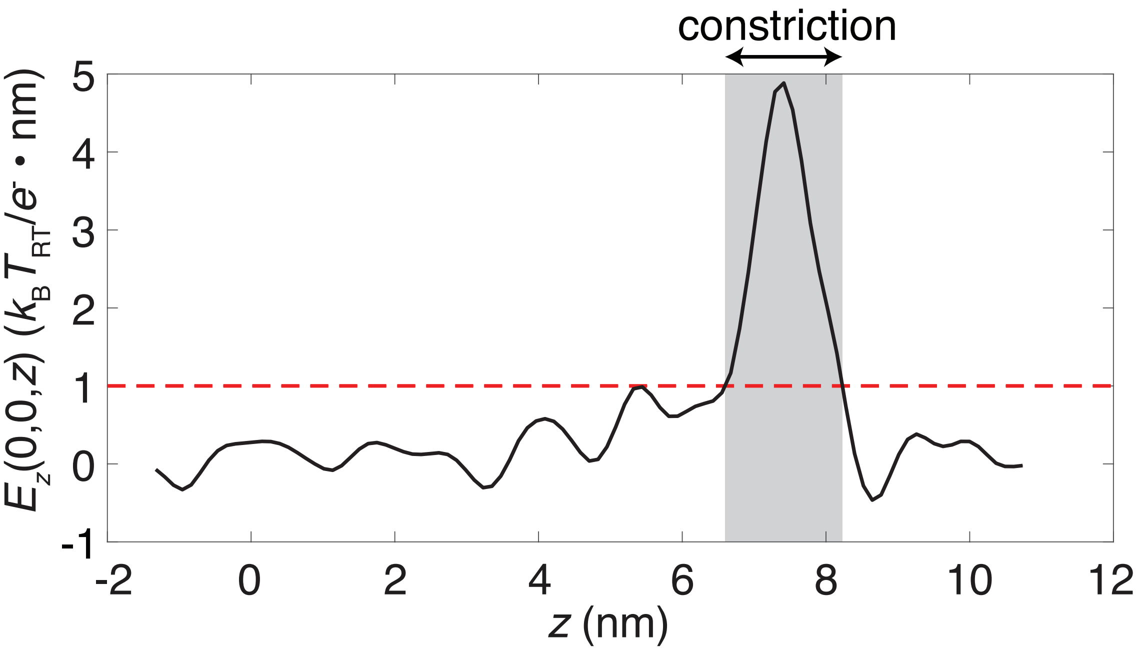


**Fig. S7** Thresholding used to define the high-field constriction of MspA. The constriction was defined as the region (shaded gray area) along the $z$-direction where the magnitude of the z-component of the electric field along the $z$-axis (computed as $E_z(0,0,z) = \nabla V_{\mathrm{ex}}(0,0,z) \cdot \hat{z}$) is greater than 1 $k_B T_{\mathrm{RT}}/e^- \cdot$nm.

Supplementary Text

### 1 Markov chain Monte Carlo calculations

To produce Fig. 3c through 3f, we used a Markov chain Monte Carlo (MCMC) calculation, also known as a Metropolis-Hastings algorithm, to produce a set of Boltzmann-distributed configurations of a model polymer confined in the MspA pore.

#### 1.1 Principle of Markov chain Monte Carlo calculations

The MCMC method is used to approximate a probability density function $p(\mathbf{x})$ given a function $f(\mathbf{x})$ (such as a Boltzmann factor $f(\mathbf{x}) = e^{-\varepsilon(\mathbf{x})/k_B T}$) such that $p(\mathbf{x}) = Af(\mathbf{x})$ where $A$ is a constant - in other words, $p(\mathbf{x})$ is proportional to $f(\mathbf{x})$. This approach is useful because in practice, $A$ is the most difficult part of $p(x)$ to compute, since normalizing $p(x)$ through straightforward integration techniques is computationally too difficult if $\mathbf{x}$ is very high-dimensional and an analytical solution cannot be found.

One such situation is polymer physics problems such as ours, since the number of degrees of freedom is $2n$, where $n$ is the number of links in the polymer chain, and we know that configurations should be Boltzmann distributed. In other words,

$$p(\mathbf{x}) = \frac{1}{\int_\Omega e^{-\beta\varepsilon(\mathbf{x}')}\,\mathrm{d}^{2n}\mathbf{x}'} \cdot e^{-\beta\varepsilon(\mathbf{x})} = A \cdot f(\mathbf{x}), \tag{1}$$

where $\beta = (k_B T)^{-1}$ and $\varepsilon(\mathbf{x})$ is the energy of a configuration given by $\mathbf{x} = (x_1, x_2, ..., x_{2n}) \in \Omega$. In this equation, the integral prefactor corresponds to the constant $A$ and the Boltzmann factor corresponds to $f(\mathbf{x})$ in the terminology used above. Because of the inclusion of numerical results for the potential in the nanopore and the shape of the hard wall defining the edge of the pore, the integral to compute $A$ yields no analytical solution. Moreover, numerically calculating $f(\mathbf{x})$ on the entirety of the domain $\Omega$, even with a relatively sparse grid, is infeasible for $n \gtrsim 5$ .

Fortunately, only knowing $f(x)$ (in our case, the Boltzmann factor), the MCMC method provides a tool for sampling from the distribution $p(x)$ in an unbiased way. A starting configuration $\mathbf{x}_0$ is chosen as well as an update distribution $\pi(\Delta\mathbf{x})$ and a number of iterations $N$. Then the algorithm proceeds as follows:

1. Initialize an iterator $i = 1$.
2. Draw a step $\Delta\mathbf{x}$ from the distribution $\pi(\Delta\mathbf{x})$.
3. Compute a candidate configuration $\mathbf{x}_{\text{new}} = \mathbf{x}_{i-1} + \Delta\mathbf{x}$.

4. Compute the Boltzmann factors $f(\mathbf{x}_{\text{new}}) = e^{-\beta\varepsilon(\mathbf{x}_{\text{new}})}$ for the candidate configuration and $f(\mathbf{x}_{i-1}) = e^{-\beta\varepsilon(\mathbf{x}_{i-1})}$ for the latest accepted configuration.

5. Compute the probability $P_{\text{accept}}$ of accepting the new configuration as the ratio of candidate to old probability densities $\frac{p(\mathbf{x}_{\text{new}})}{p(\mathbf{x}_{i-1})} = \frac{A\cdot f(\mathbf{x}_{\text{new}})}{A\cdot f(\mathbf{x}_{i-1})} = \frac{f(\mathbf{x}_{\text{new}})}{f(\mathbf{x}_{i-1})}$.

6. Accept the new configuration with probability $\max(1, P_{\text{accept}})$. If accepted, set $\mathbf{x}_i = \mathbf{x}_{\text{new}}$. Otherwise, duplicate the previous configuration by setting $\mathbf{x}_i = \mathbf{x}_{i-1}$.

After an initial "thermalization" over some number of steps $d$ during which bias from the choice of $\mathbf{x}_0$ decays, the resulting set of $N_S = N - d$ samples $S = \{\mathbf{x}_{d+1}, \mathbf{x}_{d+2}, ..., \mathbf{x}_N\}$ will be distributed according to $p(\mathbf{x})$. This sampling can then be used to estimate the value of $p(\mathbf{x})$ at different locations as

$$p(\mathbf{x}) \approx \frac{\text{number of samples } \mathbf{x}' \in S \text{ such that } \mathbf{x}' \in [\mathbf{x} - \mathrm{d}\mathbf{x}/2, \mathbf{x} + \mathrm{d}\mathbf{x}/2]}{N_S \, |\mathrm{d}\mathbf{x}|^{2n}} \tag{2}$$

where $\mathrm{d}\mathbf{x} = \frac{|\mathrm{d}\mathbf{x}|}{\sqrt{2n}}(1, 1, ..., 1)$ is a $2n$-dimensional vector defining a suitably sized box of volume $|\mathrm{d}\mathbf{x}|^{2n}$. It can also be used to find the expectation value of a function of $\mathbf{x}$,

$$\langle g \rangle = \frac{\sum_{i=d+1}^{N} g(\mathbf{x}_i)}{N_S}, \tag{3}$$

or the probability of a proposition $Q$ being true at any given moment,

$$P(Q) \approx \frac{\text{number of samples } \mathbf{x} \in S \text{ such that } Q \text{ is true}}{N_S}. \tag{4}$$

##### 1.2 Implementation for nanopore-confined polymer

To apply the MCMC method to a given problem, we only need to choose the parameters of the problem:

- The domain $\Omega$,
- the potential defining $\varepsilon(\mathbf{x})$,
- the update distribution $\pi(\Delta\mathbf{x})$,
- the inverse temperature $\beta$,
- the number of iterations $N$,
- the number of iterations to discard $d$, and
- the starting configuration $\mathbf{x}_0$.

The last four of these are simple constants. Working in units of $k_B T_{\text{RT}}$, where $T_{\text{RT}}$ is room temperature, sets $\beta = 1.136$ for our experiments at 37°C. We chose $N = 10^6$ to ensure sufficient sampling and conservatively used $d = 10^4$. We chose $\mathbf{x}_0$ to correspond to a chain fully extended downwards through the pore; how this is represented mathematically can be seen by defining the space of configurations $\Omega$.

###### 1.2.1 Modeling the DNA-peptide-DNA polymer

To model the DNA-peptide-DNA construct used in our experiments, and thereby arrive at a representation of $\mathbf{x}$ and a definition of $\Omega$, we considered a section of a variable number of chain joints separated by a distance $a_{\text{DNA}} = 0.67$ nm, followed by a section with 24 joints separated by the smaller distance $a_{\text{linker/peptide}} = 0.39$ nm, and finally another DNA section consisting of 5 joints (Fig. S5 The chain was constrained so that the first link was always at the origin, defined to be at the top of the vestibule of the pore. Charges were assigned to joints in units of electron charge depending on the identity of the monomer.

To step the chain upwards in a way analogous to Hel308, we performed $n_{\text{shift}}$ copies of the calculation, with the first DNA section being shortened each time. Since Hel308 takes approximately half-nucleotide steps of alternating size 0.55 and 0.45 nucleotides, every other step ($n_{\text{shift}} = 2, 4, 6, ...$) has the same number of joints as the polymer used in calculation $n_{\text{shift}} - 1$, while the spacing between the anchor and the first joint is shortened from $a_{\text{DNA}}$ to $0.45 a_{\text{DNA}}$.

With this model, a configuration of a polymer with $M$ joints (excluding the anchor point) can be defined as $\mathbf{x} = (\boldsymbol{\theta}, \boldsymbol{\phi}) = (\theta_1, \theta_2, ...\theta_M, \phi_1, \phi_2, ...\phi_M)$, where $\theta_i$ is the polar angle of the displacement vector pointing from joint $i-1$ to joint $i$ from the $+z$-axis, and $\phi_i$ is the azimuthal angle of that vector measured from the $+x$-axis. Thus the starting configuration $\mathbf{x}_0$ corresponding to a straight chain along the $z$-axis is simply the zero vector. This lends itself to defining an independent normal update distribution for $\theta$ and $\phi$:

$$\pi(\Delta \mathbf{x}) = (4\pi^2 \sigma_\theta \sigma_\phi)^{-M/2} \prod_{i=1}^{M} e^{-\Delta\theta_i^2/2\sigma_\theta^2} \prod_{i=1}^{M} e^{-\Delta\phi_i^2/2\sigma_\phi^2}, \tag{5}$$

where the average magnitude of perturbations made at each step to $\theta$ and $\phi$ are, in our case, defined by $\sigma_\theta = \sigma_\phi = 0.05$ radians. This choice does not affect the calculated distribution, only the time needed to adequately sample the space. If the update perturbations are too large, many candidate configurations will be rejected for intersecting with the pore wall or being energetically unfavorable. If they are too small, it will take a very large number of updates to differ significantly from the original configuration. We chose 0.05 because it corresponded to a rejection rate of roughly 50% for most constructs and shifts.

From a configuration $(\boldsymbol{\theta}, \boldsymbol{\phi})$, the Cartesian coordinates of joint $i$ can be straightforwardly computed as

$$x_i = \sum_{j=1}^{i} a_i \sin\theta_i \cos\phi_i, \tag{6}$$

$$y_i = \sum_{j=1}^{i} a_i \sin\theta_i \sin\phi_i, \tag{7}$$

$$z_i = \sum_{j=1}^{i} a_i \cos\theta_i, \tag{8}$$

where $a_i$ is the (constant) separation between joint $i-1$ and joint $i$.

###### 1.2.2 Modeling the potential energy $\varepsilon(\mathbf{x})$

We included three interactions in the potential:

1. The externally applied electric field electrophoretically trapping the DNA, resulting in an electric potential $V$ acting on the charges on the polymer,
2. Coulomb potentials between charges on the polymer, and
3. a hard wall at the surface of the nanopore.

**External field:** The externally applied potential $V_{\mathrm{ex}}$ has been previously calculated in M2MspA under a 180 mV voltage difference (Fig. S5) [2]. Since this calculation was performed on a 3D grid, we used Matlab's built-in "griddedInterpolant" function to linearly interpolate $V_{\mathrm{ex}}$ between the grid points. Points outside the domain on the *cis* side of the pore were assigned -180 mV and on the *trans* side of the pore 0 mV. The electric potential energy was thus calculated as

$$U_{\mathrm{ex}} = \sum_{i=1}^{M} q_i V_{\mathrm{ex}}(x_i, y_i, z_i). \tag{9}$$

**Polymer charge-charge interactions:** Because of the mobile ions in the salt solution, charge-charge Coulomb forces are subject to charge screening, resulting in a potential that exponentially decays with distance,

$$V_{qq} = \frac{1}{4\pi\epsilon}\frac{q\,e^{-r/\lambda_D}}{r}, \tag{10}$$

where $r$ is the distance from the a charge $q$, $\epsilon$ is the dielectric constant of the medium and $\lambda_D$ is the Debye length for the medium. For us, $\epsilon = 78\cdot\epsilon_0$ with $\epsilon_0$ being the permittivity of free space, and $\lambda_D = \sqrt{\frac{\epsilon k_B T}{2e\mathscr{M}}} = 0.40$ nm, where $\mathscr{M}$ is the ionic strength of our 400 mM KCl, 10 mM $MgCl_2$ solution. Therefore, adding up the potential energies between all pairs of charges, we obtain

$$U_{qq} = \frac{1}{4\pi\epsilon}\sum_{i=1}^{M}\sum_{j=i+1}^{M}\frac{q_i q_j\,e^{-r_{ij}/\lambda_D}}{r_{ij}}. \tag{11}$$

**Exclusion from the pore wall:** The boundary of the interior of the pore was drawn by taking a cross-section of the potential map $V_{\text{ex}}$ and drawing the minimal boundary that excluded the high-potential regions inside the protein or bilayer (Fig. S5). The boundary was drawn by choosing points by hand, through which a spline $r_p(z)$ was computed defining the radius of the pore $r_p$ as a continuous function of $z$ position. Outside of the vertical boundaries of the pore, $-1.14$ nm $\leq z \leq 8.95$ nm, $r_p$ was set to infinity. To prevent any transgressions of the polymer through the pore wall, we implemented a hard infinite potential wall at this boundary:

$$U_{\text{wall}} = \begin{cases} \infty & \text{if } x_i^2 + y_i^2 \geq r_p(z_i)^2 \text{ for any } i \\ 0 & \text{otherwise.} \end{cases} \tag{12}$$

The total potential energy then is $\varepsilon = U_{\text{ex}} + U_{qq} + U_{\text{wall}}$.

#### 1.3 Analysis of Markov chain Monte Carlo results

Fig. 3f was produced by using equation 4 with respect to the presence of a phosphorylated amino acid in the constriction defined by 6.6 nm $\leq z \leq$ 8.2 nm. These boundaries were determined by thresholding the $z$-component of the potential gradient down the $z$-axis to find the region of highest field in the nanopore (Fig. S7, also see Fig. S5 for an annotation of the constriction over the MspA pore radius and applied potential), where charges present most affect the ion current.

If $z_{\text{Ph}}$ is the $z$-coordinate of the joint corresponding to the phosphorylated amino acid, then for the two constructs with a single phosphorylation we computed the "phosphorylation weight" $W_{\text{Ph}}$ as

$$W_{\text{Ph}} = \frac{\text{\# of configurations where } 6.6 \text{ nm } \le z_{\text{Ph}} \le 8.2 \text{ nm}}{N_S} \tag{13}$$

while for the construct with two phosphorylations at $z$-positions $z_{\text{A}}$ and $z_{\text{B}}$ we simply computed

$$W_{\text{Ph}} = W_{\text{Ph,A}} + W_{\text{Ph,B}}. \tag{14}$$

To generate the video S1, we created a 2-D histogram for each shift with even bins in $r = \sqrt{x^2 + y^2}$ and in $z$. Each bin of the histogram was populated with a count for every sample in the MCMC results where the phosphorylated joint fell into that bin, as per equation 2, and normalized by the volume of the bin to obtain a density. Since the bin volume is not rectangular but instead a ring shape, as we are marginalizing over the $\phi$ coordinate, the volume is $2\pi r \cdot \Delta r \Delta z$, where $\Delta r$ and $\Delta z$ are the bin spacings in $r$ and $z$.

(1) Derrington, I. M., et al. Subangstrom single-molecule measurements of motor proteins using a nanopore. Nature biotechnology 33.10, 1073-1075 (2015).

(2) Bhattacharya, S., et al. Water mediates recognition of DNA sequence via ionic current blockade in a biological nanopore. ACS nano, 10(4), 4644-4651 (2016).
